# Repetitive Mild Head Trauma Induces Activity-Mediated Lifelong Brain Deficits in a Novel *Drosophila* Model

**DOI:** 10.1101/2021.02.09.430429

**Authors:** Joseph A. Behnke, Changtian Ye, Aayush Setty, Kenneth H. Moberg, James Q. Zheng

**Author notes:** **Corresponding Author**: Dr. James Q. Zheng, Department of Cell Biology, Emory University School of Medicine, Atlanta, GA 30322.

## Abstract

Mild head trauma, including concussion, can lead to chronic brain dysfunction and degeneration but the underlying mechanisms remain poorly understood. Here, we developed a novel head impact system to investigate the long-term effects of mild head trauma on brain structure and function, as well as the underlying mechanisms in *Drosophila melanogaster*. We find that *Drosophila* subjected to repetitive head impacts develop long-term deficits, including impaired startle-induced climbing, progressive brain degeneration, and shortened lifespan, all of which are substantially exacerbated in female flies. Interestingly, head impacts elicit an elevation in neuronal activity and its acute suppression abrogates the detrimental effects in female flies. Together, our findings validate *Drosophila* as a suitable model system for investigating the long-term effects of mild head trauma, suggest an increased vulnerability in brain injury in female flies, and indicate that early altered neuronal excitability may be a key mechanism linking mild brain trauma to chronic degeneration.

## Introduction

Repetitive head trauma, including mild injuries such as concussion, is a risk factor for developing several neurodegenerative disorders, including Alzheimer’s disease (AD) [1,2], Parkinson’s disease (PD) [3], Amyotrophic lateral sclerosis (ALS) [4] and chronic traumatic encephalopathy (CTE) [5–9]. Although each of these conditions arise from distinct etiologies, they share some common features including abnormal protein deposits [3, 4, 10, 11], impaired axonal transport [12], swelling [13] and retraction [14], immune cell activation, and excitotoxicity [15–22], which ultimately result in progressive cellular degeneration, cell death, and frank brain atrophy. Symptomatic onset typically appears later in life in each of these aforementioned diseases, yet neurodegeneration is believed to begin years in advance [23, 24]. While repetitive head trauma is a risk factor for developing neurodegeneration, it remains a challenge to provide a causal link between the mechanisms underlying the early injury response and chronic cellular and behavioral sequelae. A more complete understanding of the brain’s response to injury over time will lend new insight towards the development of neuroprotective strategies.

The use of rodent models is a common method to study the basic pathophysiological mechanisms of traumatic brain injury [16, 25–32], but the relatively long rodent lifespan together with the use of surgical procedures for delivering injuries [27, 33] represent a challenge to investigate the onset and progression of lifelong effects following head trauma. Here, we developed a tractable *Drosophila melanogaster* (fruit fly) model that enables lifelong interrogation of specific processes and their downstream cellular and behavioral effects. Fruit flies represent an excellent tractable model organism to dissect fundamental cellular and molecular disease mechanisms, including neurodegeneration [34–36], while taking advantage of their relatively short lifespan [37–42] that enables long-term outcome analyses related to head injury exposure. Several fly models have been developed to study traumatic brain injury in *Drosophila*, but the lack of controlled head-specific impacts may confound neuronal-specific responses [43–45]. Head-specific impact models have recently been developed in which physical head impacts were delivered to the heads of individually restrained flies [46–48], but no long-term study was carried out to examine the detrimental effects that might develop late in life.

In this study, we devised a novel injury model with high precision and accuracy in delivering repetitive headfirst impacts to a large number (10-15 flies at a time) of awake and unrestrained fruit flies. We then applied this model to investigate novel mechanisms related to repetitive mild head impacts on lifelong function in both sexes. We established that early exposure to repetitive mild head impacts resulted in long-term behavioral dysfunction and brain degeneration, which is more pronounced in female flies. We further investigated an underlying mechanism and identified neuronal activity as a key mediator for mild trauma-induced long-term brain dysfunction. Together, this work provides a causal relationship between the early phase of injury involving neuronal activity and the subsequent chronic degenerative processes.

## Materials and Methods

### Fly husbandry

Flies were maintained at 18°C (activity-suppressing experiments) or 25°C, with 60% humidity on a 12-hr:12-hr light:dark cycle and kept in vials containing fresh fly media consisting of cornmeal, yeast, molasses, agar, p-hydroxy-benzoic acid methyl ester. Vials were changed every 3–4 days using sterile methods. The following stocks were used: *w^1118^* (BDSC 5905), *UAS-CaLexA-LUC*[49] (from K. Abruzzi), *OK107-GAL4* (BDSC 854), *neuronal synaptobrevin-GAL4 (nSyb-GAL4)[50]* (from T. Ngo), *nSyb-LexA*[50] (from T. Ngo), *LexAop-Shibire^ts1^* [50] (from T. Ngo), *Oregon R* (BDSC 5). A stable line expressing *nSyb>Shibire^ts1^ (LexA>LexAop)* was created using the respective aforementioned stocks and used for pan-neuronal activity suppressing experiments. Unless otherwise noted, flies used were progeny from *nSyb-LexA* males crossed to female *w^1118^*. For activity suppressing experiments, injured flies were progeny from *nSyb>Shibire^ts1^* stably-expressing males crossed to female *w^1118^*.

### High-speed video recording and Video-Assisted Startle-Induced Climbing Assay

High-speed video recording of head impacts was acquired using a Phantom Miro M310 high-speed camera controlled by a PC with Phantom Camera Control (PCC) software. Sequences recorded at either 3205 or 4106 frames per second were imported into Image J. Body orientation at impact was measured using the Angle tool.

Startle-induced climbing was assessed using a modified negative geotaxis assay. Flies were placed in vials containing 5% agar. Up to four vials were assayed at the same time, using a customized 3-D Polylactic Acid (PLA) printed rig. For each trial, the rig was lightly tamped three times, and fly movement was recorded using a Panasonic HC-V800 digital video recorder at 60 frames per second. Individual vials from each video were cropped, and the first 10s were trimmed in Image J and underwent automated fly behavior tracking using id.tracker.ai. All testing took place between ZT 3 and ZT 8 (ZT, Zeitgeber time; lights are turned on at ZT 0 and turned off at ZT 12) and testing occurred between 20-22°C under normal lighting conditions.

### Automated Fly Behavior Tracking using idtracker.ai

Automated fly behavior was performed using idtracker.ai, a deep-learning algorithm and software that permits individual tracking of flies. All videos were processed on a Google Cloud Compute Virtual Machine (VM) running a PyTorch Deep Learning VM Instance (c2-deeplearning-pytorch-1-2-cu100-20191005) equipped with an N1-Standard-16 (16 vCPUs, 60 GB memory) machine type, an NVIDIA Tesla T4 GPU, a 100GB SSD disk and Intel Haswell CPU platform. Subsequent data analysis was performed using custom Matlab scripts.

### Immunohistochemistry for Detecting Neurodegeneration

Whole flies were fixed for 3 hours in 4% paraformaldehyde (PFA) in phosphate-buffered saline containing 0.5% Triton-X (PBS-T). The flies were rinsed 4 times for 15 minutes each with PBS-T and brains were subsequently dissected in PBS-T. Brains were permeabilized overnight in 0.5% PBST at 4 °C with nutation then blocked in 5% normal goat serum (NGS) in PBS-T for 90 minutes at room temperature with nutation. Brains were stained with DAPI (1:1000; Invitrogen D1306) (nuclear marker) and Alexa Fluor 594 (AF594) phalloidin (1:400, Invitrogen A12381), which binds to filamentous actin and stains the whole-brain parenchyma). Regions devoid of DAPI and phalloidin are considered vacuoles (Video S4). Stained brains were rinsed 4 × 15 minutes with PBS-T, followed by one wash in PBS for 90 minutes and then mounted on glass slides within SecureSeal Imaging Spacers (Grace Bio-Labs, Bend, Oregon, USA) containing SlowFade Gold Antifade mounting medium (Life Technologies, Carlsbad, CA, USA S36937). Whole-brain imaging was performed using a two-photon FV1000 laser-scanning confocal (Olympus) to acquire 1 μM thick sections. Image analysis was performed using ImageJ (Fiji) software.

To demonstrate that vacuoles detected using this strategy correspond to neuropil loss, additional confirmatory staining was performed on a subset of brains for pre- and post-synaptic markers (mouse monoclonal α-bruchpilot [1:30; DSHB nc-82] and mouse monoclonal α-discs large 1 [1:50; DSHB 4F3], respectively) (Fig. S1). The secondary antibody was AF-568 labeled goat anti-mouse Ig (1:400; Invitrogen A11031). Alexa Fluor 488 (AF488) phalloidin (1:400, Invitrogen A12379) replaced the use of AF594 phalloidin. Whole-brain imaging of 1 μM thick sections was performed using either a two-photon FV1000 laser-scanning confocal (Olympus) or a Nikon C2 laser-scanning confocal.

### Neuronal Activity Luciferase Reporter

To measure luciferase activity, representative groups of 2-4 *OK107>CaLexA- LUC* sham and injured flies were quickly immobilized using CO2, decapitated and heads collected in 100μL 1x Promega Glo Lysis Buffer (Promega #E2661), and 4-7 independent samples were collected for each treatment group. Fly heads were homogenized using an automated pestle motor mixer (Argos Cat. No. A0001) for 1 minute at room temperature, then incubated at room temperature for 1h, centrifuged for 10 min at 13.5k RPM. The supernatant was transferred to a new tube. For luciferase assays, 60 μL of each sample was mixed 1:1 with Steady-Glo Luciferase system (Promega #E2510) following the manufacturer’s protocols, and then added to a white-walled 96-well plate at room temperature and incubated in the dark for 10 minutes. Luminescence was measured using a BioTek Synergy HTX Multi-Mode Reader.

### Data Reporting and Statistical Analysis

No statistical methods were used to determine sample sizes but are consistent with sample sizes similar to those generally employed in the field [51]. Experimenters were not blinded as almost all data acquisition and analysis was automated. All flies in each vial were administered with the same treatment regimen. For each experiment, the experimental and control flies were collected, treated, and tested at the same time. A Mann-Whitney U test with Bonferroni or Holm correction for multiple comparisons was used for statistical analysis of behavioral data. All statistical analyses were performed using R (R v3.5, [rstatix]). P values are indicated as follows: **** P < 0.0001; *** P < 0.001; ** P < 0.01; and * P < 0.05. For boxplots, lower and upper whiskers represent 1.5 interquartile range of the lower and upper quartiles, respectively; boxes indicate lower quartile, median, and upper quartile, from bottom to top. When all points are shown, whiskers represent range and boxes indicate lower quartile, median, and upper quartile, from bottom to top.

## Results

### Development of a novel repetitive head impact *Drosophila* model

To investigate the long-term consequences of repetitive head trauma, we developed a novel repetitive head injury model using *Drosophila* that accurately and precisely delivers headfirst acceleration-deceleration-mediated head impacts (Fig. 1a, Video S1). Multiple flies (10-15 flies at a time) are contained within a customized plastic injury vial designed to fit within our injury rig consisting of a cradle that is connected to a pulley system with a specific counterweight. Each injury trial consists of pulling the cradle downward to the base of the injury rig, followed by lightly tamping the cradle three times for all the flies to fall to the bottom of the vial before releasing it for upward acceleration. When the cradle, which contains the injury vial with flies suddenly stops at the top, the momentum of the fruit flies continues to propel them upward, where they sustain headfirst impacts against the glass ceiling at the top of the vial (Fig. 1a, Video S1). The light tamping is employed for two purposes: to cause all the flies to fall to the bottom so they will travel the same distance before the head impact and to allow the flies to orientate in a head-upward posture as a part of their startle-induced negative geotactic climbing response. Indeed, our high-speed video recording revealed that a majority of the flies hit the top ceiling with their heads first, resulting in an impact angle of nearly 90 degrees (anterior-posterior body axis perpendicular to the impact surface, see Fig. 1a). This tailored approach improves upon an existing strategy that delivers traumatic injury to flies within a vial that is accelerated in a circular motion [44], which causes the flies to undergo a tumbling motion, thereby prompting subsequent nonspecific impacts sustained to systemic regions of the body other than the head (Fig. 1b) and limiting the head-specific conclusions that can be drawn from this type of injury. With our current impact configuration, we determined using the high-speed video recording that each fly experienced an impact that involves ~1 ms of contact at a speed of ~4 meters/s with the glass ceiling (Fig. 1a, arrows indicate immediately before and after contact between the fly and impact surface; asterisks indicate contact with impact surface). Immediately following headfirst impact, flies fell to the floor of the vial and exhibited a concussive-like behavioral response indicative of neurological injury (Fig. 1c, Video S2) whose frequency increases with the number of successive impacts incurred; these behavioral signs include: temporary loss of consciousness and other uncoordinated behavioral responses (Video S2). Importantly, flies sustain no gross morphological damage to the head or body following repetitive head impact exposure, indicative of a closed-head injury (Fig. S2a).

**Fig 1.**
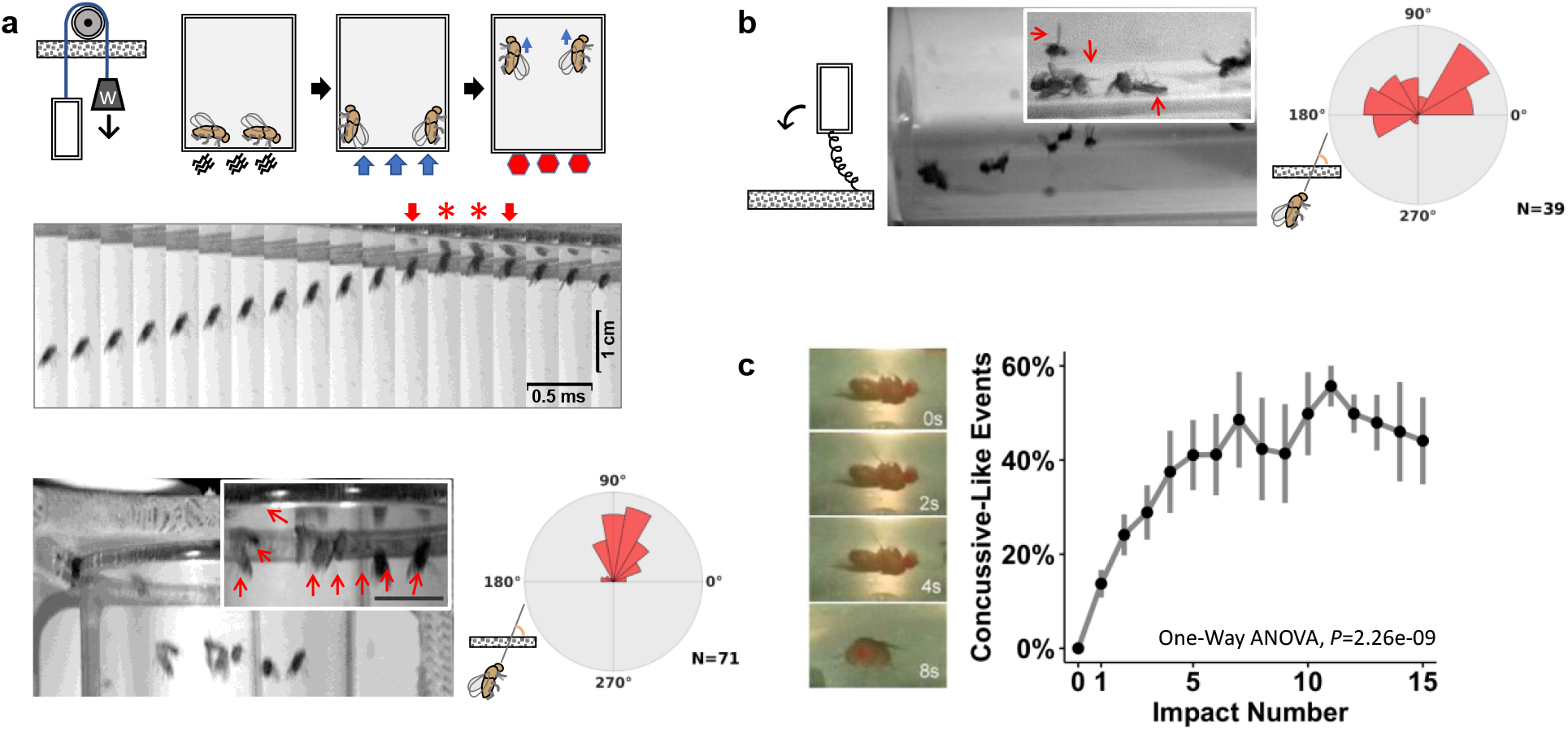
Development of a novel repetitive head impact *Drosophila* model. **(a)** *(Top)* Diagram of our novel headfirst impact model that utilizes a counterweight pulley system to accelerate a vial of flies upward ultimately resulting in acceleration-deceleration headfirst impacts. To prime proper body orientation, the vial is lightly tamped down, resulting in the natural negative geotactic upwards orientation, after which the vial is release, causing the counterweight to accelerate the vial unidirectionally. The unidirectional movement upwards, together with the priming sequences encourages facilitates headfirst orientation at impact once the vial reaches its maximum height. *(Middle)* High-speed image sequence showing headfirst movement through impact with the surface of the injury vial (arrows indicate frame immediately before and after contact, which is denoted by red stars, which lasts ~1ms long). *(Bottom)* Angular histogram comparisons demonstrate more reliable and consistent headfirst impacts and improves upon **(b)** previously existing strategies which encourage rotational body movement that results in inconsistent head orientation at impact. **(c)** Immediately following headfirst impacts, flies sustain acute signs of neurological injury, such as temporary loss of consciousness that becomes more prevalent with increasing number of repetitive head impacts. Impact number increases concussive-like events (one-way ANOVA, F (1,89)=44.26645, *P=*2.26e-09). Data represented as mean ± SD, n=4-8 vials (10 flies per vial) of both male and female flies.

### Acute recovery of climbing deficits following minimally lethal repetitive head impacts is sexually dimorphic

To emulate repetitive mild head injury exposure sustained across time, young adult (3 days post-eclosed) male and female flies were subjected to two sessions of repetitive impacts 24h apart; each session consisting of 15 iterative impacts delivered 10s apart (Fig. 2a). Repetitively impacted flies exhibited minimal (<7%) acute mortality following both days of impacts (Fig. 2b), whereas a single head impact generated no acute death (Fig. S2b). This degree of mortality is far less than the mortality (20-60%) seen in a commonly employed method used to deliver repetitive head trauma to flies[44, 52], indicating that our approach delivered a much milder insult (thus considered as “mild head trauma”). Following the injury, flies were rested for 1.5h before functional recovery testing was performed. This length of recovery time enabled acute recovery from temporary uncoordinated and seizure-like behaviors seen shortly following injury [48], and allowed flies to regain postural stability needed to perform the behavioral function assay. To measure acute behavioral deficits following repetitive head impacts, startle-induced climbing was assessed using the negative geotaxis assay (NGA) [53–56], which is a widely used locomotor assay sensitive to age-related decline and neurodegeneration. This assay tests both vestibulomotor and sensorimotor functions which are commonly assessed in traditional mammalian TBI models [57–59]. Flies were placed in plastic vials containing 2mL of 5% agar at the bottom and subjected to gentle tamping to induce a startle response. Videos were analyzed using an automated python-based tracking software, *idtracker.ai* [60, 61], to measure cumulative climbing distance (slope), total (final) and vertical distance traversed in a 10s span of time following the startle stimulus (Video S3). Testing was performed 1.5h after the first and second sessions of repetitive impacts (“1.5h” and “25.5h” with respect to the initial injury), as well as 24h after each session to assess longer recovery (“24h” and “48h” with respect to the initial injury). While injured males showed an initial deficit in climbing after each session that resolved 24h later, female flies exhibited a progressive climbing deficit that was further exacerbated following the second session of impacts (Fig. 2c-d, Fig. S3a&c). 24h after the second session, a robust difference in climbing behavior was seen between injured sexes, with injured female flies showing a 30% reduction in climbing whereas injured males performed similar to their sham counterparts (Fig. 2e).

**Fig 2.**
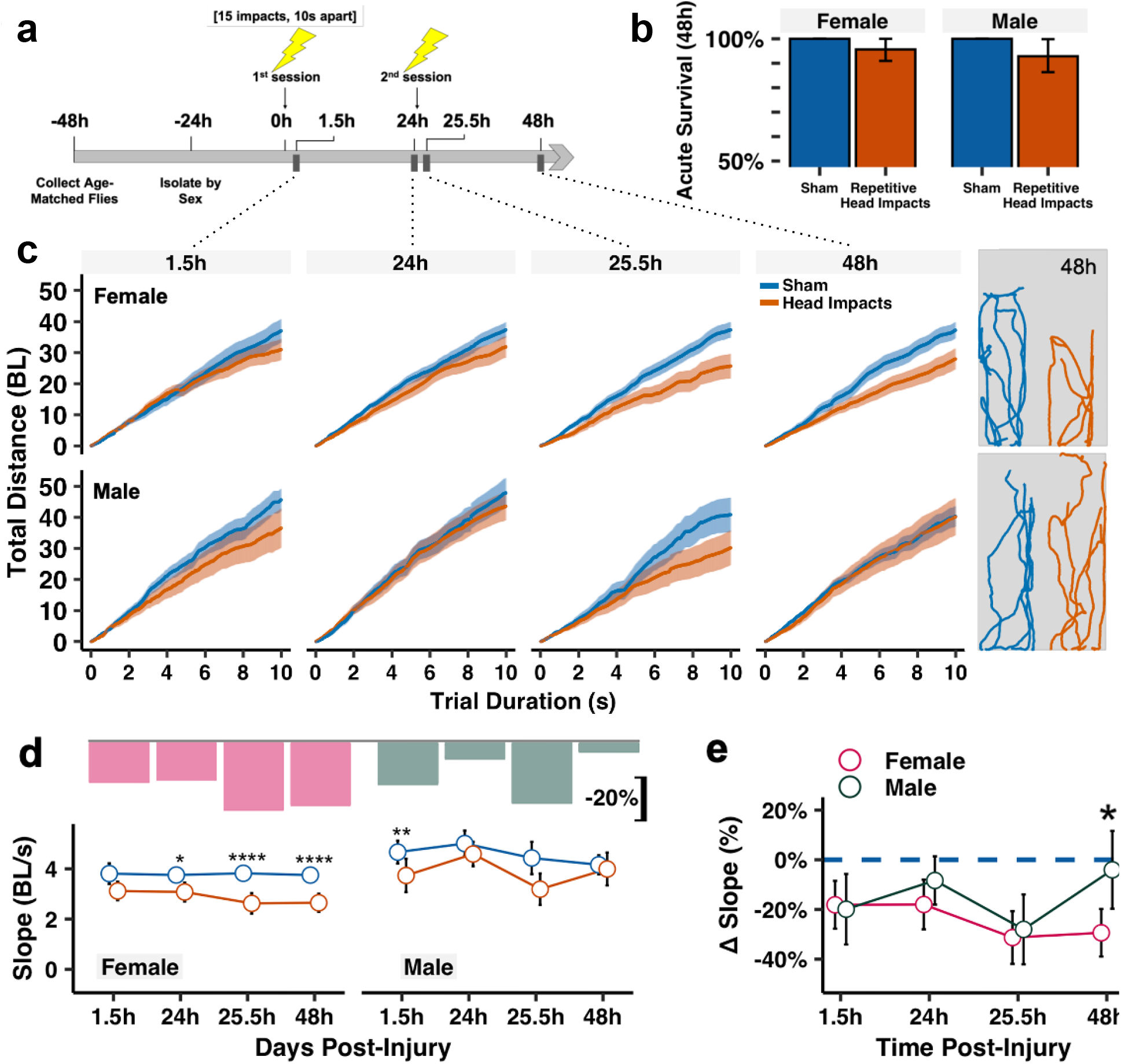
Acute recovery of climbing deficits following minimally lethal repetitive head impacts is sexually dimorphic. **(a)** Injury timeline schematic showing that flies are subjected to two sessions of repetitive head impacts (24h apart). Each session consists of 15 iterative impacts spaced 10s apart. Behavior and mortality are longitudinally monitored throughout life. **(b)** Barplot of acute survival following repetitive impacts with 95% confidence intervals. n=56-69 flies per injured group/sex and n=42-50 flies per sham group/sex. **(c)** Total cumulative distance traversed during the negative geotaxis assay. Plotted values are median with 95% confidence intervals as shaded regions, *(Right)* Representative movement tracings from 5 representative flies/group of sham (blue) and injured (orange) flies during the 10s trial duration at 48h. Distance units are in body lengths (BL). **(d-e)** Plotted climbing slope showing that injured females exhibit a progressive decrease that worsens after the second impact session, while injured male flies show active acute recovery 24h after each session of impacts. Bar plots in **(d**) correspond to the median relative decrease in climbing performance between injured and sham groups (**Δ** slope= (Injured Slope-Median Sham Slope)/Median Sham Slope). Plotted values are median with 95% confidence interval error bars in **(d-e)**. Mann–Whitney *U* test between **(d)** injured and non-injured groups, with Holm correction and **e** injured female and male performance relative to non-injured, with Bonferroni correction. *p<0.05, **p<0.01, ***p<0.001, ****p<0.0001, n=25-35 flies per sex/time/injury group.

### Lifelong deficits after early exposure to repetitive mild head impacts

One key advantage of our *Drosophila* model of mild head trauma is that it enables the monitoring and examination of long-term effects on lifespan and brain structure and function in relatively large cohorts. We thus continued to routinely examine flies following repetitive mild head impacts over their lifespan. We first measured the mortality and found that repetitive head impact exposure resulted in a shortened overall lifespan that was more prominent in female flies (Fig. 3a&b). To determine the effect of repetitive head injury exposure on long-term behavioral function, startle-induced climbing performance was routinely assessed until 98 days (14 weeks) post-injury. Flies subjected to repetitive head injury exposure exhibited long-term climbing deficits that were more profound in female than male flies (Fig. 3c-e, Fig. S4). Significant deficits in climbing performance of injured female flies were consistently observed up to 70 days post injury (Fig. 3c-d). During this time period, injured female flies exhibited a greater reduction in climbing ability compared to injured males (Fig. 3e). By 84 days post-injury, female flies from the sham and injury groups exhibited no apparent difference in their climbing performance, which is likely attributed to the severely reduced baseline performance due to aging. On the other hand, male files appear to perform better at these later ages (84- and 98-days), as such, a small difference between injured and sham male flies in climbing performance existed up to 98 days post-injury (Fig. 3c-d, Fig. S4). Together, this data demonstrated that early exposure to repetitive mild head trauma disrupts motor functions that lasts over a long period of time and disproportionally impacts female flies.

**Fig 3.**
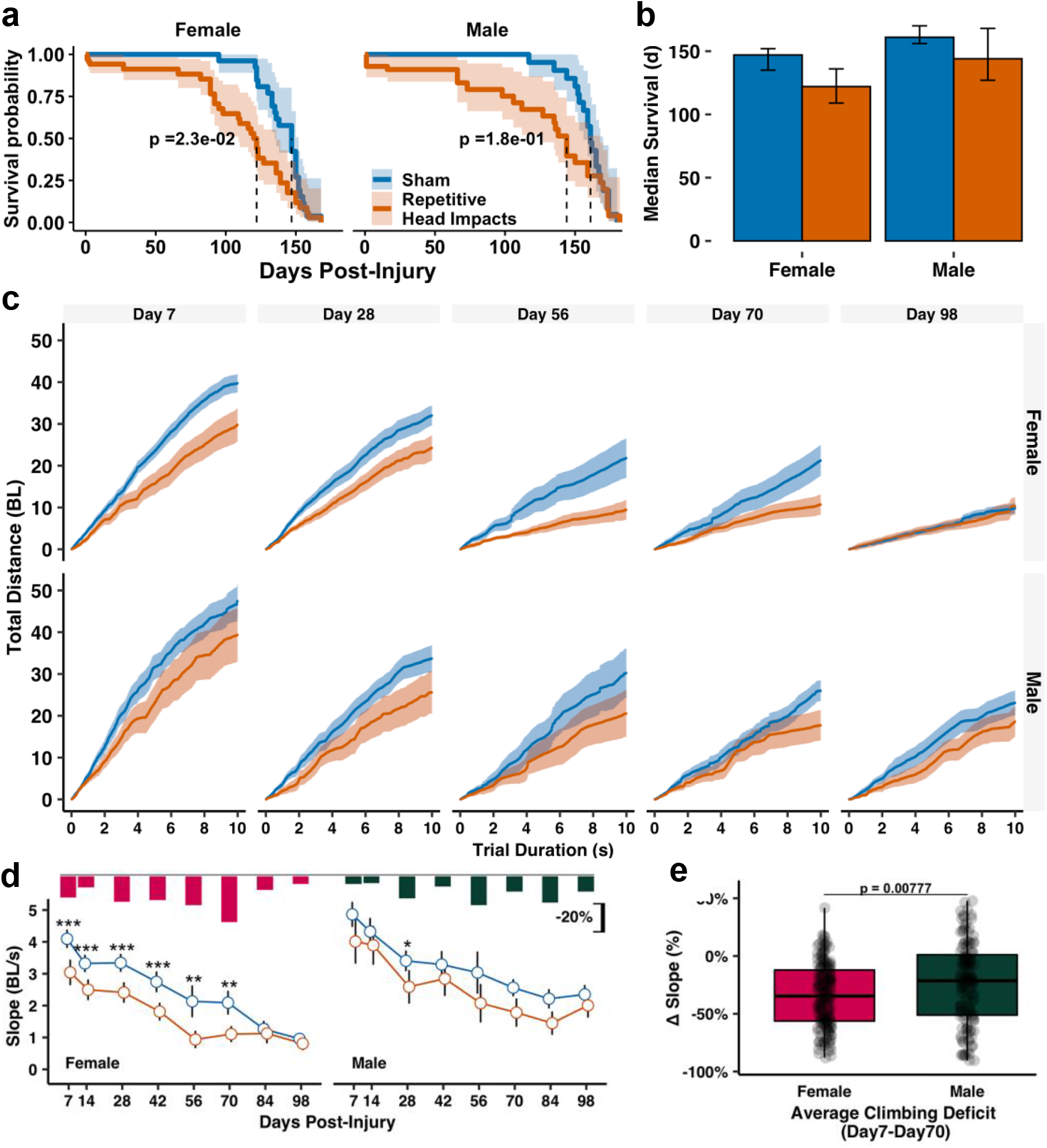
Repetitive head impacts result in a shortened lifespan and long-term behavioral climbing deficits. **(a,b)** Repetitive head impacts result in a shortened overall lifespan that is significantly different in injured female flies compared to sham injured flies. Kaplan-Meier *p*-values were determined using the Mantel-Cox log rank test with Bonferroni correction. **(c-e)** Repetitive head impacts exacerbate age-related climbing deficits that are more pronounced in female flies. Plotted values are median with 95% confidence intervals as shaded regions in **(c)** or error bars in **(b&d**) Bar plots in **(d)** indicate median relative decrease in climbing performance between injured and sham groups (**Δ** slope= (Injured Slope-Median Sham Slope)/Median Sham Slope). Boxplots in **(e)** contain individually plotted **Δ** slope values with whiskers corresponding to the maximum 1.5 interquartile range. Mann-Whitney *U* test between **(d-e)** injured and non-injured groups, with Holm correction: *p<0.05, **p<0.01, ***p<0.001, n≥22 flies per sex/time/injury group except day 56 (n≥11 flies).

### Repetitive head impacts exacerbate age-related brain degeneration

To examine brain pathology related to repetitive head injury exposure, whole-brain gross neurodegeneration was longitudinally assessed by measuring vacuolization; a commonly used measure of neurodegeneration in *Drosophila* [40, 62]. To measure vacuolization, whole-brains were stained with phalloidin to detect actin-rich neuropil and DAPI to detect nuclei and imaged using two-photon microscopy. Regions devoid of both DAPI and phalloidin signals were characterized as vacuoles [42]. Brains from flies subjected to repetitive impacts were processed for neurodegeneration acutely (1.5h) and chronically (5 weeks) post-injury. Although minimal to no vacuolization existed at the early time point across sexes and injury groups (Fig. S5), aged brains exhibited a far greater degree of vacuolization, as measured by the number and area of vacuoles per brain (Fig. 4). Both female and male flies subjected to repetitive head impacts resulted in an increase in neurodegeneration five weeks post-injury relative to sham (Fig. 4a-d), seen as an increase in the overall number of vacuoles per brain (Fig. 4e) and overall area of vacuoles per brain (Fig. 4f).

**Fig 4.**
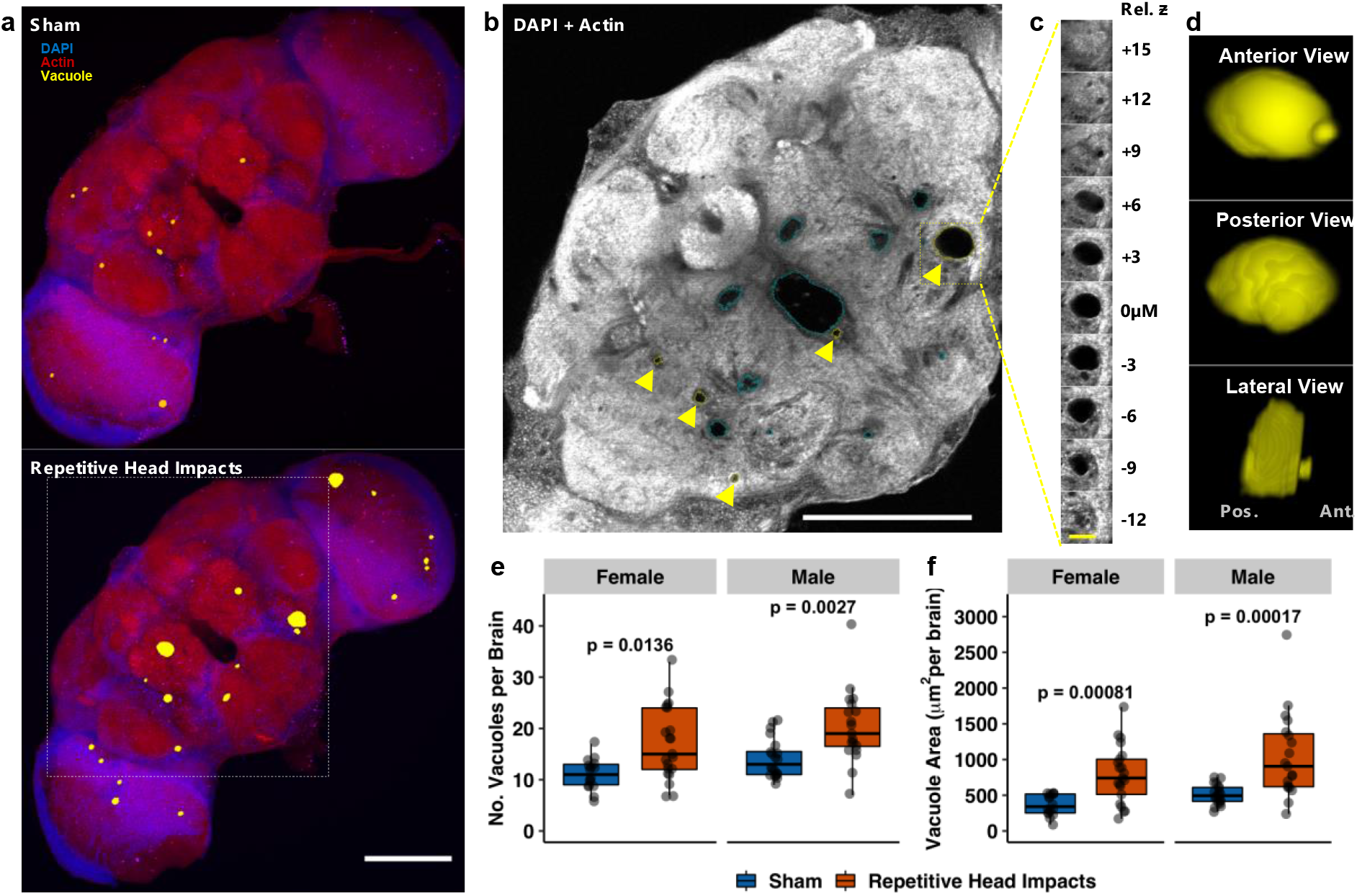
Repetitive head impacts exacerbate age-related neurodegeneration five weeks after injury. **(a)** Representative max projection of whole-brain slices (1μM thick) imaged using two-photon microscopy with brain vacuoles (neurodegeneration) depicted as yellow overlays which correspond to regions devoid of DAPI (blue) and phalloidin (red) signal from a *(Top)* sham injury brain and *(Bottom)* repetitively injured brain, captured five weeks following repetitive head impact exposure (2 sessions of 15 impacts). **(b)** Representative two-photon microscopy slice from white square outline of injured brain in **a** where pathological vacuoles of varying size (enclosed within yellow outline with yellow arrow) are found throughout the midbrain while blue outlines designate physiologically normal holes. **(c-d)** Representative vacuole outlined in **(b)** is depicted in **(c)** as a series of z-stack images and **(d)** 3-D reconstruction of the vacuole. **(e,f)** Quantification of **(e)** vacuole number and **(f)** vacuole area per brain from sham and repetitively injured brains. Boxplots contain individually plotted values with whiskers corresponding to the maximum 1.5 interquartile range. Within sex differences between sham and repetitive head impact conditions were analyzed with the Mann–Whitney *U* test with Bonferroni correction. White scale bar=100μM, yellow scale bar=20μM.

### Elevated brain activity by repetitive mild head impacts and the protective effects by acute suppression of neuronal activity are sex-dependent

We next began to explore the mechanisms underlying the development of long-term behavioral and brain dysfunction after repetitive mild head impacts. Emerging evidence indicates that neuronal hyperexcitability plays an important role in a number of neurodegenerative disorders [63]. Neuronal hyperexcitability has been examined in moderate to severe TBI [16, 17], but its existence in mild TBI remains poorly understood. We thus investigated if repetitive mild head trauma exposure in *Drosophila* elicits neuronal activity *in vivo*. To do so, we measured neuronal activity using a non- invasive *in vivo* luciferase-based transcriptional indicator of calcium signaling, *CaLexA- Luciferase (CaLexA-LUC)* [49, 64] (**<u>Ca</u>**lcium-dependent nuclear import of **<u>LexA</u>**); a Nuclear factor of activated T-cells (NFAT)-LexA chimera whose transcriptional activity is based on the Ca^2+^-regulated dephosphorylation and nuclear translocation of NFAT which then drives luciferase expression. Pan-neuronal expression of UAS-CaLexA using the neuronal synaptobrevin-GAL4 *(nSyb-GAL4)* is weak and sparse [51], and in our hands, results in sick and sluggish appearing flies (data not shown). Instead, we measured neuronal activity in an essential midbrain structure, the mushroom body [65], using the robust mushroom body driver to express CaLexA *(OK107>CaLexA-LUC)*. The mushroom body contains roughly 2% of the total fly brain neurons [65] and features a dense organization of projection neurons that communicate with various regions of the fly brain, making it a center for high-level integration of functions related to learning/memory [65] and startle-induced climbing [66]. *OK107>CaLexA-LUC* flies were subjected to repetitive impacts (1 session of 30 impacts) and collected for fly head lysate preparation 1.5 hours following injury. Bioluminescence (luciferase activity) was measured from collected fly head lysates. Injured female flies exhibited a 20% increase in measured neuronal activity relative to their sham female counterparts, whereas no significant difference was seen between injured and sham male flies (Fig. 5a).

**Fig 5.**
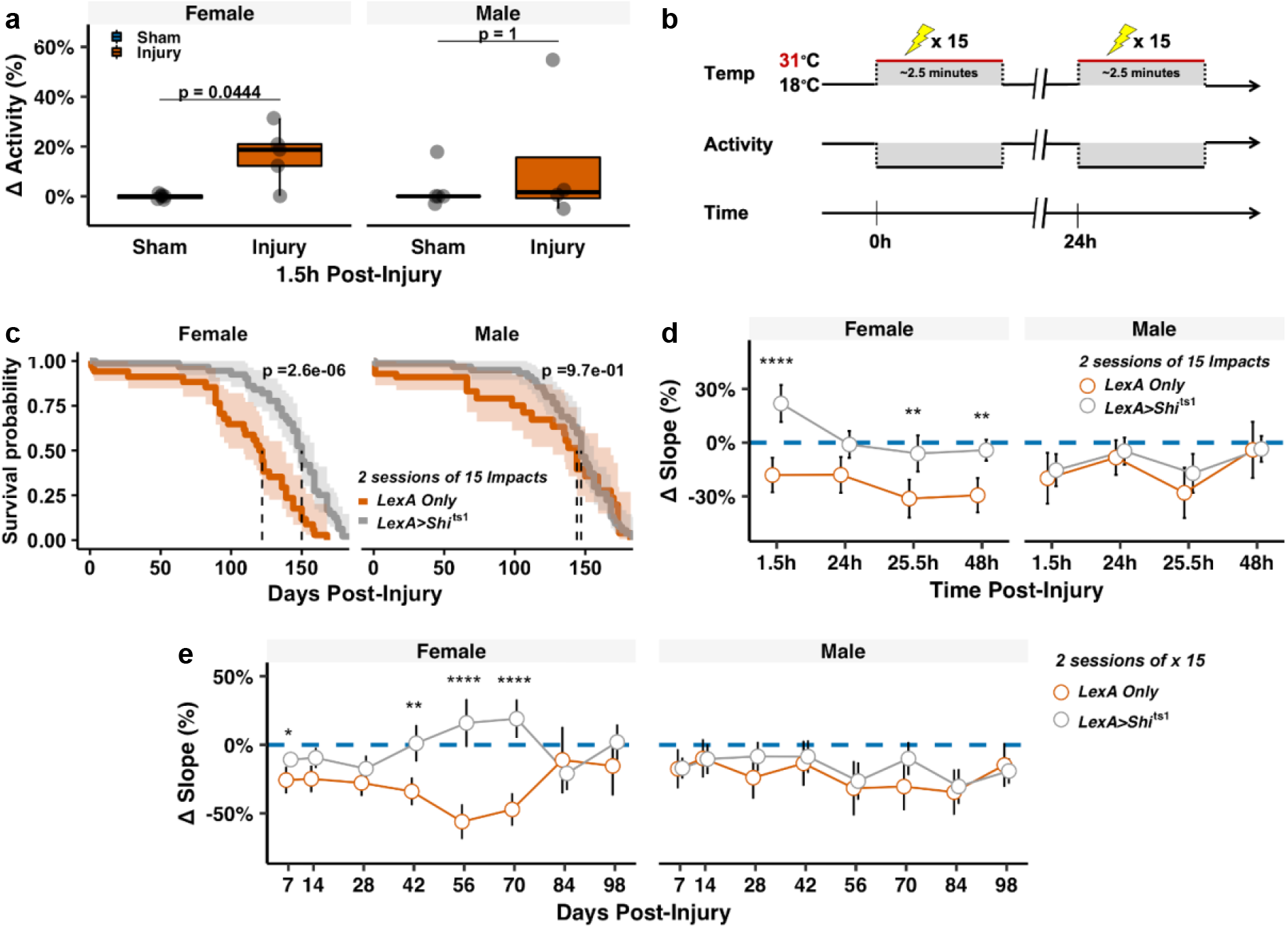
Long-term benefit of suppressing acute injury-induced neuronal activity following repetitive head impacts preferentially affects females. **(a)** Quantification of neuronal activity using mushroom body *in vivo* calcium monitoring (*OK107>CaLexA- LUC)* revealed an acute increase in neuronal activity 1.5h following repetitive head impact exposure in lysates collected from injured female brains, but not males. Within sex differences between injury and sham were analyzed using the Mann-Whitney *U* test with Bonferroni correction, n=4-7 lysates per group (2-4 fly heads per lysate). **Δ** activity= ((Sample luminescence)- (Median Sham Luminescence))/(Median Sham Luminescence). Boxplots contain individually plotted values with whiskers corresponding to the maximum 1.5 interquartile range. **(b)** Schematic overview of strategy to conditionally silence neuronal activity using the pan-neuronally expressed temperature-sensitive hypomorphic null allele, *Shibire^ts1^* during the repetitive injury induction period. 18°C → 31 °C shift conditionally suppresses neuronal activity for 2.5 minutes during delivery of repetitive head impacts in pan-neuronally *shi^ts1^*-expressing flies *(LexA>shi^ts1^)*. **(c)** Kaplan-Meier survival curves showing that blocking neuronal activity at the time of injury protects against shortened lifespan following injury exposure which preferentially benefits female flies. *p*-values were calculated using the Mantel-Cox log rank test with Bonferroni correction and correspond to within sex differences between injured *LexA only* and *Shibire^ts1^-containing* flies *(LexA>shi^ts1^)*. **(d,e)** Blocking activity protects against **(d)** acute and **(e)** chronic climbing deficits related to repetitive head impact exposure that are preferentially found in females. Plotted values in **(d-e)** are relative median slopes (relative to respective uninjured sham) with 95% confidence interval error bars. Differences in climbing behavior were analyzed using the Mann– Whitney *U* test with Holm correction, between injured *LexA Only* and *Shibire^ts1^*-containing flies *(LexA>shi^ts1^)*. *p<0.05, **p<0.01, ***p<0.001, ****p<0.0001.

We then took a tailored approach to conditionally suppress neuronal activity at the time of injury, using the temperature sensitive dynamin homologue mutant, *shibire^ts1^* (*shi^ts1^*) [50, 67, 68] expressed pan-neuronally (*nSyb>shi^ts1^*). Pan-neuronal expression of shi^*ts1*^ enables conditional suppression of synaptic activity by halting synaptic vesicle release, via interruption of the readily releasable pool of synaptic vesicles, thereby inducing a reversible state of flaccid paralysis [50, 67, 68]. Conditional suppression of synaptic activity occurs rapidly when exposed to the non-permissive temperature threshold of 30°C, which is initially achieved through transfer to a pre-warmed vial and maintained via exogenous heat delivered by a heat lamp.

Control flies expressing either pan-neuronal *LexA* only *(nSyb-LexA)* or pan-neuronal *shibire^ts1^ (nSyb>shi^ts^)* flies were injured during two sessions of 15 impacts delivered 24h apart. During each injury session, flies were subjected to the non-permissive temperature for only the extent of the injury induction period (~2.5 minutes long) (Fig. 5b). Suppression of neuronal activity at the time of injury nearly abolished the acute mortality found in both injured *nSyb-LexA* flies and restored normal female lifespan (Fig. 5c). Suppression of neuronal activity at the time of injury restored relative acute female climbing performance compared to *nSyb-LexA* injured performance, whereas injured *nSyb>shi^ts1^* males showed no significant improvement compared to injured *nSyb-LexA* (Fig. 5d; Fig. S6). Moreover, long-term climbing performance of injured *nSyb>shi^ts1^* females better resembled their sham *nSyb>shi^ts1^* counterparts compared to injured *nSyb-LexA* flies (Fig. 5e; Fig. S7). This finding was most prominent during Days 42-70 post-injury. Interestingly, the long-term relative performance of injured male *nSyb>shi^ts1^* flies appeared no different than injured *nSyb-LexA* flies, which may be due to the milder initial climbing deficit seen in injured males (Fig. 3).

We next examined if blocking acute activity at the time of injury is protective against long-term neurodegeneration following repetitive head impacts. Similarly, repetitive head impacts were delivered to *nSyb>shi^ts1^* flies at the non-permissive temperature and injured *nSyb>shi^ts1^* flies were processed using the same methods shown in Fig. 4 for vacuole assessment following the aforementioned acute-silencing regimen. Acute silencing of activity at the time of injury was found to substantially reduce the number and size of vacuoles in female brains, while no obvious reduction in injury-induced vacuole formation was found in injured male brains from *nSyb>shi^ts1^* compared to *nSyb-LexA* (Fig. 6). Together with the mortality and climbing data, this data suggests that acute aberrant neuronal activity elicited by repetitive head trauma may potentiate long-lasting dysfunction that preferentially affects female flies. In accordance with this finding, blocking acute activity through the expression of the dominant negative transgene, *shibire^ts1^* provides long-term neuroprotection against repetitive injuries that preferentially benefits females.

**Fig 6.**
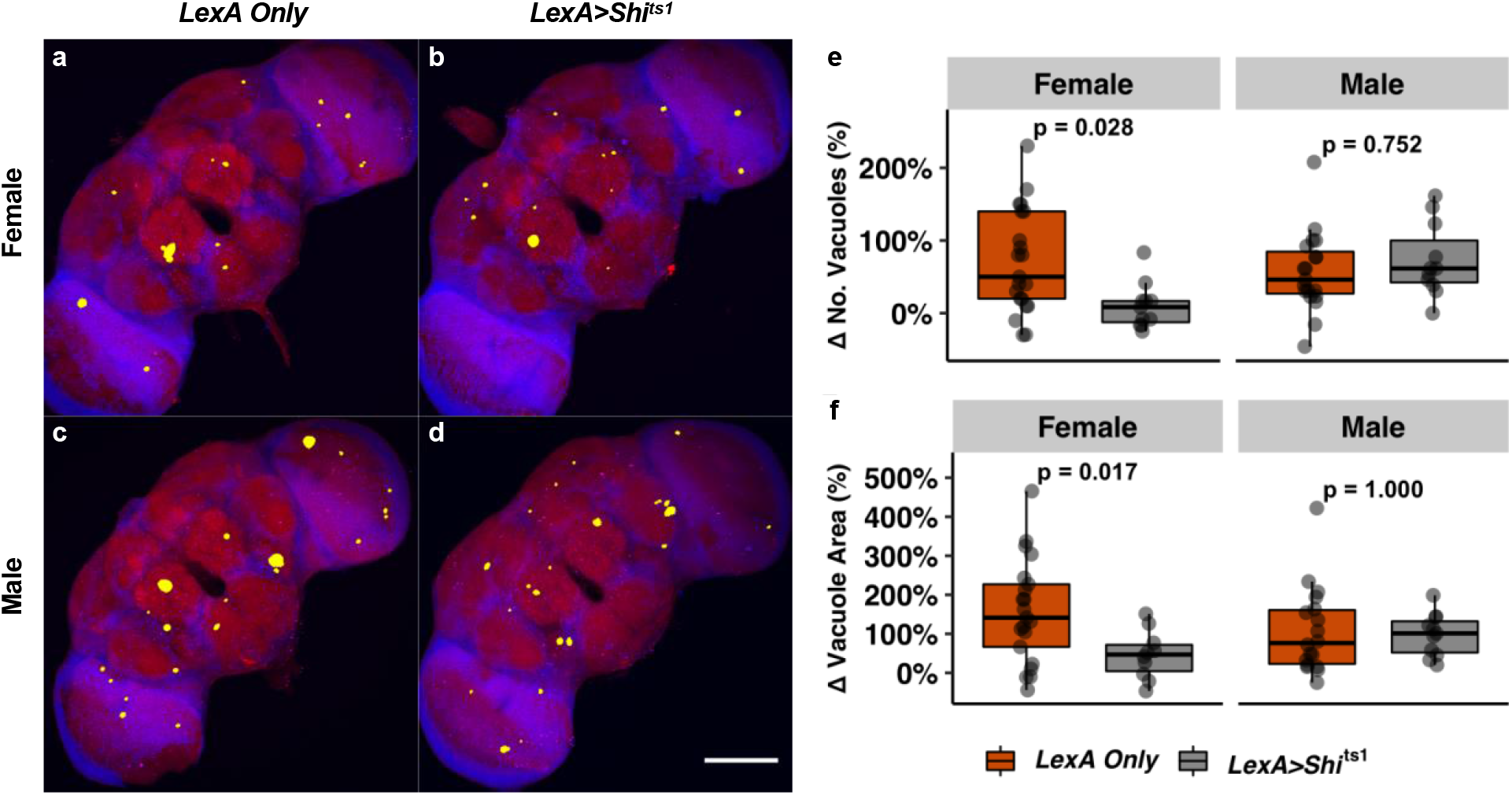
Acute suppression of neuronal activity mitigates injury-induced chronic neurodegeneration in a sex-dependent manner. **(a-d)** Representative max projection of whole-brain slices (1μM thick) imaged using two-photon microscopy with brain vacuoles (neurodegeneration) depicted as yellow overlays which correspond to regions devoid of DAPI (blue) and phalloidin (red) signal from injured **(a)** female *nSyb-LexA* brain, **(b)** female *nSyb>Shi^ts1^* brain, **(c)** male *nSyb-LexA* brain, and **(d)** male *nSyb>Shi^ts1^* brain. White scale bar=100μM. **(e, f)** Boxplot of relative quantification of **(e)** vacuole number and **(f)** vacuole area (relative to respective uninjured sham) with whiskers corresponding to the maximum 1.5 interquartile range. Within sex differences between injured *nSyb-LexA (LexA Only)* and *nSyb>Shi^ts1^ (LexA>Shi^ts1^)* were analyzed with the Mann–Whitney *U* test with Bonferroni correction. White scale bar=100μM.

## Discussion

An emerging body of evidence demonstrates that a past history of repetitive mild head trauma increases the risk for developing cognitive and motor impairment, and brain degeneration later in life [5, 6]. However, it remains unclear how the early exposure to mild head trauma leads to the development of long-term neurological dysfunction. This is in part due to the challenges using existing animal models for lifelong examination and interrogation of the detrimental effects after injury. In this study, we developed a *Drosophila* head impact system to investigate the long-term detrimental effects of mild repetitive head trauma and their underlying mechanisms. Our data show that repetitive mild physical impacts delivered to the adult fly head result in acute concussion-like responses and impairments in startle-induced climbing behavior. Importantly, when monitored over an extended period of time (throughout the entire lifespan), the early repetitive head trauma was found to shorten lifespan and elicit a decline in behavior, particularly in female flies. Whole-brain imaging also revealed increased brain degeneration in flies that received early repetitive head trauma. Finally, we provided evidence that neuronal activity plays a critical role in mild head trauma-induced long-term deficits in behavior and brain pathology within female flies.

While fruit flies are anatomically different from human, it has become clear that *Drosophila* is an excellent model organism to investigate the molecular, cellular, and genetic mechanisms underlying a wide range of human disorders. For example, the use of *Drosophila* to has led to the discovery of novel genes and mechanisms of neurodegeneration [36, 69–73], including those involved in axonal transection-mediated degeneration, that has informed and has been validated by subsequent murine [74–76] and human studies [77]. The findings from our study not only support the use of *Drosophila* as a model system to study the long-term brain degeneration and dysfunction developed by environmental insults, but also provide crucial insight into the underlying mechanisms involving excitatory neuronal activity in the development of brain deficits induced by head trauma. Our finding that silencing neuronal activity largely abolishes the long-term detrimental effects on fly behavior and brain pathology is consistent with the emerging view on the involvement of excitotoxicity in a number of neurodegenerative disorders [63] including TBI [78–80]. Lastly, our findings also suggest that lifelong injury outcomes are exacerbated in the female flies compared to male flies.

Our newly developed *Drosophila* head impact model possesses several unique features that distinguish it from other systems. First, the physical impacts are delivered mostly to the head of awake and unrestrained flies. The headfirst collision of flies to the impact ceiling is achieved in part through the upward body orientation at the start of acceleration due to the innate negative geotactic escape response during which flies orient headfirst against the force of gravity when startled. The aerodynamic body shape of the flies is also believed to contribute to the headfirst impact orientation as we observed mostly headfirst impacts with flies in a coma (data not shown). Second, the headfirst impacts are delivered to a large cohort of flies (10-15 flies at a time), thus increasing the efficiency for investigating the effects on large populations that are essential for analyzing genetic variations and long-term age-dependent changes, the latter of which is made possible by the relatively short overall lifespan of fruit flies. This, together with our non-invasive automated behavioral testing paradigm, is amenable to future high-throughput screening for genetic and environmental modifiers. Third, injury severity, or the impact force sustained by the fly head can be specifically modified by adjusting the hardness of the impact surface. For example, our current glass impact surface, which is a circular glass coverslip, can be replaced with a material surface of different hardness, such as plastic or Styrofoam, which can provide a specific degree of buffering for mimicking the padding of helmets in contact sports. Injury severity can also be adjusted within our system by using different counterweights and heights (for various acceleration distances) within the pulley system, resulting in a change in velocity at impact. Therefore, our head impact system has the potential to be utilized in a wide range of studies concerning head trauma that are focused on behavioral, structural, and functional consequences using adult flies of different ages and genetic backgrounds, helping to better understand the risk factors in the development of neurodegeneration and long-term dysfunction after head trauma.

While most affected individuals appear to recover from mild head trauma, the risk for developing long-term deficits in brain function and structure has been documented [5]. One interesting finding is that our data revealed that female flies were preferentially more affected by repetitive head impacts than males. This was seen behaviorally, as a greater reduction in climbing ability relative to sham injured flies, as well as a shortened overall lifespan. The greater persistent functional deficit found in female flies following injury parallels human data from the Transforming Research and Clinical Knowledge in Traumatic Brain Injury (TRACK-TBI) study in which the female sex was associated with decreased six-month functional outcome measured using the Glasgow Outcome Scale-Extended (GOSE) [81]. A recent cross-sectional human study revealed that females were more likely to report a less favorable health-related quality of life (HRQoL) during the chronic stage of TBI (10 years post-injury) [82]. Additional evidence demonstrates that young female athletes may take longer to become symptom free following sports-related concussion [83]. Taken together, this demonstrates sex differences following recovery from TBI and substantiates the importance of using both sexes in preclinical animal models. Interestingly, sex-dependent differences in function (climbing behavior) in fly head injury models has only been examined once prior, in which no differences between sex were revealed [43]. However, flies were only measured for 72 hours during the post-injury period, and the method for measuring behavior was done using the traditional approach to measure startle-induced climbing performance [53, 54]. To provide greater sensitivity to potential subtle differences in climbing behavior, we used idtracker.ai [60], an automated deep-learning tracking algorithm that enables measurement of individual fly performance. Climbing function was measured until 98 days post-injury, which accounted for most of the overall lifespan for flies reared at 18°C (~150 day median survival).

At this moment, the underlying mechanism for the greater vulnerability to mild head trauma for females remains unclear. It may be related to females being more likely to report mild TBI [84]. There are also a number of biological factors that may account for sex-dependent differences as well. Gender differences in head-neck anatomy result in greater head acceleration (angular acceleration and displacement) in females compared to males [85, 86], which may elicit a greater force acted upon the head and thus increase the injury risk found in females[86]. Female flies are known to have a relatively larger body size compared to males [87, 88], which may result in a greater impact force and worse outcomes following injury. Future high-speed camera studies are needed to address this consideration in greater detail.

Sexually dimorphic vulnerability to injury may in part be attributed to neuroanatomical differences, which have been shown to exist *in vitro* [89]. In response to mechanical trauma, sudden stretching of the axon can increase membrane permeability, resulting in the temporary loss of ionic homeostasis, an influx of cations and excessive excitatory neurotransmitter release, and subsequent excitotoxicity, metabolic dysfunction and cell death [15–19]. Cultured neurons from female human and murine sources feature axons with fewer microtubules per axon, resulting in a thinner axon diameter compared to males [89], making it potentially more vulnerable to trauma. Moreover, in response to injury using a cell stretch injury model, female axons exhibited a greater acute influx of calcium compared to males [89], which may explain the finding of more persistent deficits in hippocampal synaptic plasticity following mild TBI, seen in preclinical data from female rats [90]. Our study provides corresponding *in vivo* data demonstrating that repetitive injury elicits acute neuronal activity within the injured female fly brain. Although this surge of excitation is thought to be short-lived, on the scale of minutes to hours [91,92], this action may potentiate long-lasting effects on synaptic plasticity and neuronal function and affect the brain’s vulnerability to further injury, however this theory remains untested in part to the lack of animal models for long-term examination.

Here in this study, we were able to target neuronal activity, specifically neurotransmitter release, at the time of injury using the transgenic temperature-sensitive *shibire^ts1^* mutant fly line as a tool to conditionally suppress whole-brain neuronal activity and monitor mortality, lifelong functional deficits and brain degeneration. Our data provides the first evidence that silencing neuronal activity at the time of injury was able to largely mitigate the long-term detrimental effects of mild repetitive brain trauma in female flies. Together, this supports a mechanism involving synaptic plasticity and neuronal excitability related to chronic trauma-induced degeneration. Future work using our system can assess the potential benefit of silencing activity at various timepoints following repetitive injuries to determine the long-term therapeutic window of active rest, which is a commonly prescribed treatment following head injury. Additional work will also longitudinally examine how early repetitive head impact exposure affects neuronal excitability later in life, especially given that increased neuronal excitability is associated with brain aging [93], for which injury exposure may accelerate.

Although blocking activity seemed to preferentially benefit injured females to a greater extent than injured males, this may be related to the timing and duration of the activity suppression. Within our current paradigm, neuronal activity suppression was limited to the time of the injury exposure (~2.5 minutes), after which flies were quickly returned to the permissive temperature for the remainder of their lives. Additional strategies could be used within our current system that provide chronic suppression of changes in hyperexcitability using glial cells involved in excitatory neurotransmitter recycling by specifically targeting excitatory neurotransmitter transporters [94–96]. The inability to properly regulate excitability may be related to the loss of excitatory neurotransmitter buffering capacity in glial cells, primarily astrocytes, which have been shown to be perturbed in severe isolated scenarios of head trauma [97, 98]. While this remains to be seen in repetitive mild head trauma exposure, decreased buffering capacity may potentiate chronic changes in hyperexcitability that then result in eventual neuronal degeneration. Interestingly, the early stages of Alzheimer’s disease may involve abnormal changes in glutamatergic buffering, which results in persistent excessive neuronal activation, excitotoxicity and eventual cell death [99, 100]. Activity-dependent changes related to repetitive exposure to trauma may represent an early mechanism that potentiates AD, as well as other neurodegenerative diseases. Additional studies using our tractable injury model can also examine the role of hyperexcitability towards the formation of abnormal protein deposition seen in AD and CTE[8, 11, 101].

## Conclusions

This work utilizes a novel *Drosophila* head impact model to investigate how early exposure to repetitive mild head trauma results in long-term brain dysfunction and degeneration. We find that exposure to repetitive trauma resulted in lifelong brain dysfunction and degeneration that is further exacerbated in female flies. We also uncovered a novel activity-dependent mechanism underlying head trauma-induced long-term brain deficits. The finding that silencing neuronal activity during head impacts can effectively mitigate the development of long-term deficits suggests it as a potential therapeutic target for treating chronic sequalae of repetitive mild head trauma exposure. The basis of this work calls to question whether treating *milder*, potentially unassuming injuries, should receive greater attention as a means to thwart long-term complications. Understanding the natural course and lifelong implications of head trauma exposure is critically important for developing long-term neuroprotective strategies. We will continue to use our tractable fly model to investigate genetic and environmental factors that affect neurodegeneration secondary to repetitive head trauma exposure and provide insight into the fundamental mechanisms that lead to the progression of lifelong neurological dysfunction.

## Authors’ contributions

J.A.B., and J.Q.Z. designed research; J.A.B. and C.Y. performed research; J.A.B., C.Y., A.Y. and J.Q.Z. analyzed data; J.A.B. and J.Q.Z. designed and built the head injury device; J.Q.Z. and K.H.M. supervised research; and J.A.B., C.Y. and J.Q.Z. wrote the paper.

## Acknowledgements

We thank Drs. Janet Alder, Smita Thakker-Varia and Michael Sayegh for their comments on the manuscript. We also thank Dr. Brian Robinson and Chris Rounds for their technical assistance with *Drosophila* handling, and Dr. Lauren Stewart at Georgia Institute of Technology for her help with high-speed video recording. Corey Zheng at Georgia Institute of Technology helped with the design and construction of the Pulley impact system as well as the 3-D printing of the impact cradle. Thank you to Teri Ngo (Rubin Lab) and Dr. Kate Abruzzi (Rosbash Lab) for providing fly stocks. This research project would not have taken shape without stimulating discussions with Drs. Dorothy Lerit and Victor Faundez, and other colleagues within the Department of Cell Biology at Emory University.

