## Supplemental Data for "Repetitive Mild Head Trauma Induces Activity-Mediated Lifelong Brain Deficits in a Novel *Drosophila* Model"

**Supplemental Data (Behnke et al.)**
Part 1: Legends for Videos S1-S4
Part 2: Figures S1-S7

### Part 1: Legends for Videos S1-S4

**Video S1 Demonstration of headfirst impacts using our novel *Drosophila* model** Multiple unrestrained awake flies contained within a plastic injury vial are accelerated upward until the vial reaches the top of the apparatus, after which forward momentum of the flies carries them further upward until they impact the upper surface of the vial where they sustain headfirst impacts

**Video S2 Head impacts produce immediate concussive-like behavior** Flies sustain acute signs of neurological injury immediately following head impacts, such as temporary loss of consciousness and uncoordinated behaviors that become more prevalent with increasing number of repetitive head impacts.

**Video S3 Automated tracking of fly climbing using idtracker.ai** Individual fly climbing behavior within the startle-induced negative geotaxis assay is analyzed using an automated tracking algorithm idtracker.ai.

**Video S4 Novel approach to measure frank neurodegeneration in *Drosophila* whole-brain mounts** Two-photon microscopy stack of whole-brain mount stained for DAPI (nuclei) and phalloidin (actin for brain parenchyma). Regions devoid of DAPI/phalloidin that are shaded in green correspond to physiologically normal holes while red shaded regions correspond to pathological vacuoles.

**Part 2: Figures S1-S7**

**Figure S1**

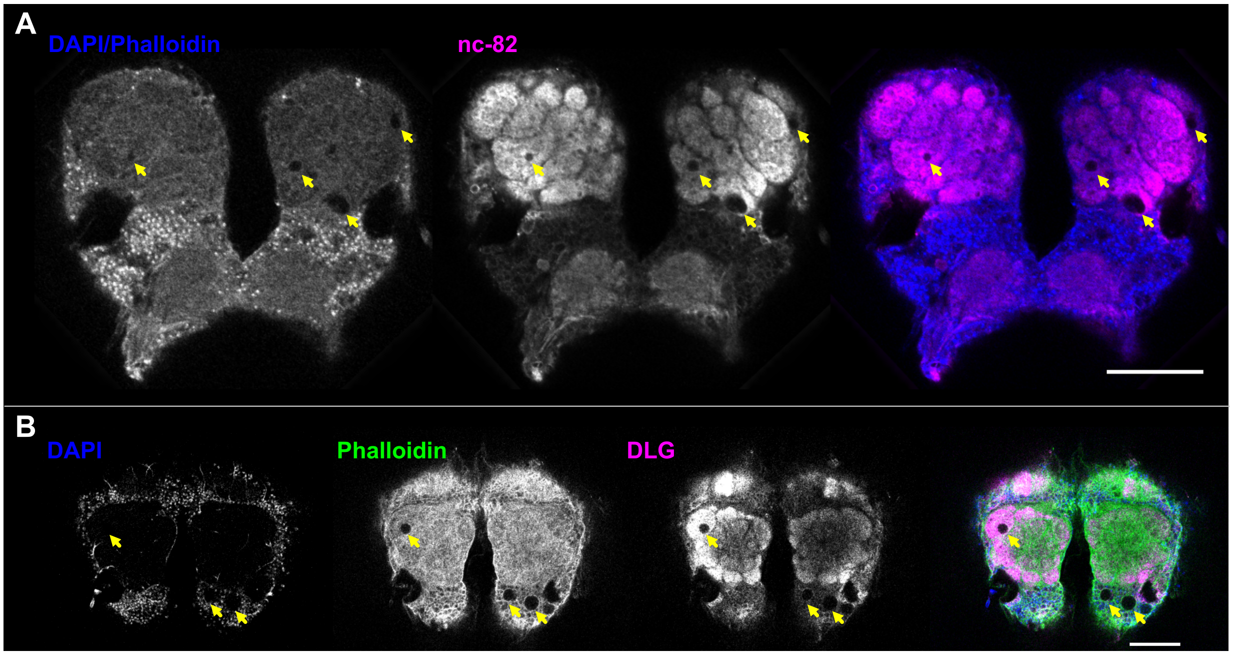


**Detecting Neurodegeneration in Drosophila Whole-Brain Mounts** (**a**) Two-Photon Microscopy of whole-brain mounts stained with DAPI and phalloidin (blue) to detect brain parenchyma and nc-82 (bruchpilot, magenta) to detect pre-synaptic neuropil. (**b**) Confocal Microscopy of whole-brain mounts stained with DAPI (blue) to detect nuclei, phalloidin (green) to detect brain parenchyma, and discs large 1 (DLG, magenta) to detect post-synaptic neuropil. Yellow arrows designate vacuoles (absence of signal), scale bar = 50μm.

**Figure S2**

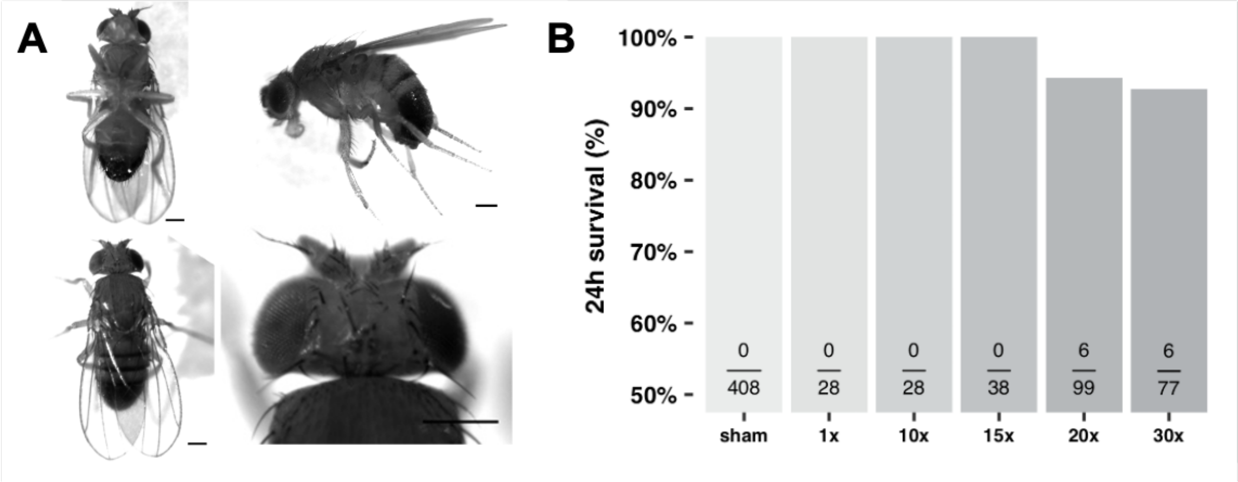


**Gross Morphology and Acute Survival Following Repetitive Head Impacts (a)** Representative whole-body micrograph of an injured *Oregon R* male fly showing no signs of gross morphological damage to the head or body following repetitive head impact exposure. Scale bar= 100 µm**. (b)** Barplot of acute survival (24h) following varying number of iterative successive head impacts delivered 10s apart. Black text indicates (# dead/# at risk).

**Figure S3**

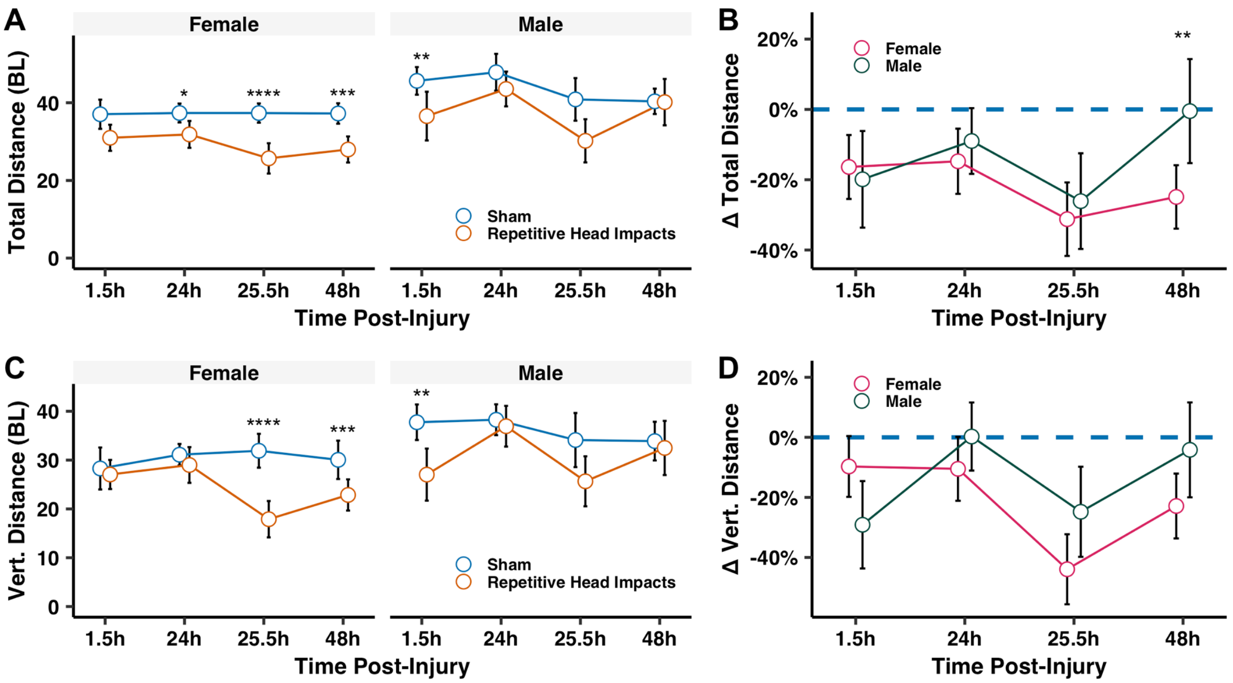


**Acute recovery of climbing deficits following minimally lethal repetitive head impacts is sexually dimorphic.** Repetitive head impacts elicit acute climbing deficits in both male and female, seen as a reduction in **(a)** total climbing distance and **(c)** total vertical distance traversed during the startle-induced climbing assay. **(b&d)** Injured female flies show progressive relative behavioral deficits that worsen after the second impact session, while injured male flies show active acute recovery 24h after each session of impact. Plotted values are median **(a&c)** raw or **(b&d)** relative (to respective sex) values with 95% confidence intervals as error bars. Mann–Whitney *U* test between **(a&c)** injured and non-injured groups, with Holm correction and **(b&d)** injured female and male performance relative to non-injured, with Bonferroni correction. *p<0.05, **p<0.01, ***p<0.001, ****p<0.0001, n=25-35 flies per sex/time/injury group.

### Figure S4

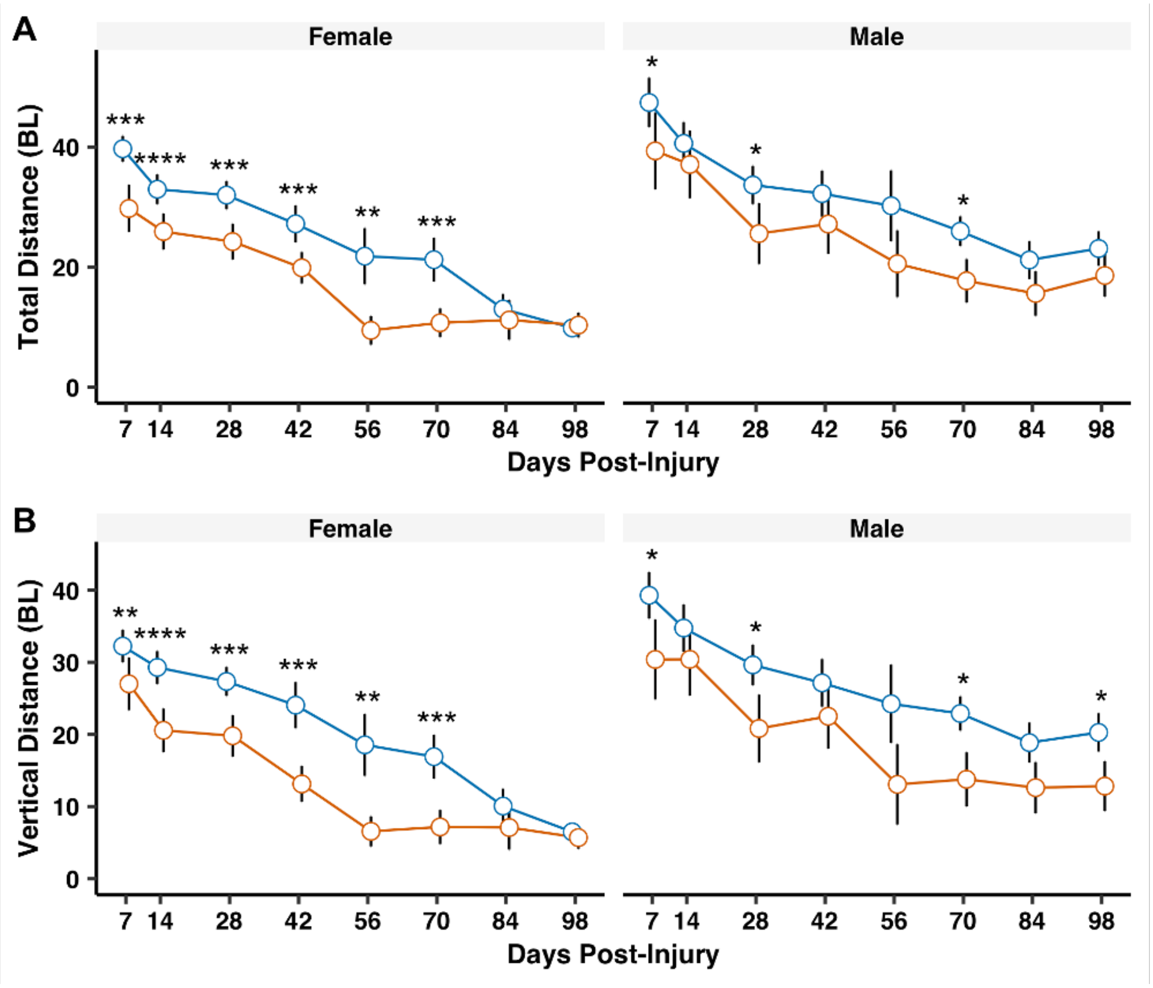


**Repetitive head impacts result in long-term behavioral climbing deficits.** Repetitive head impacts elicit chronic climbing deficits that are more pronounced in female flies, seen as a reduction in **(a)** total climbing distance and **(b)** total vertical climbing distance traversed during the climbing assay. Plotted values are median values with 95% confidence intervals as error bars. Mann–Whitney *U* test between injured and non-injured groups, with Holm correction: *$p<0.05$, **$p<0.01$, ***$p<0.001$, ****$p<0.0001$, $n\geq22$ flies per sex/time/injury group except day 56 $n\geq11$ flies.

**Figure S5**

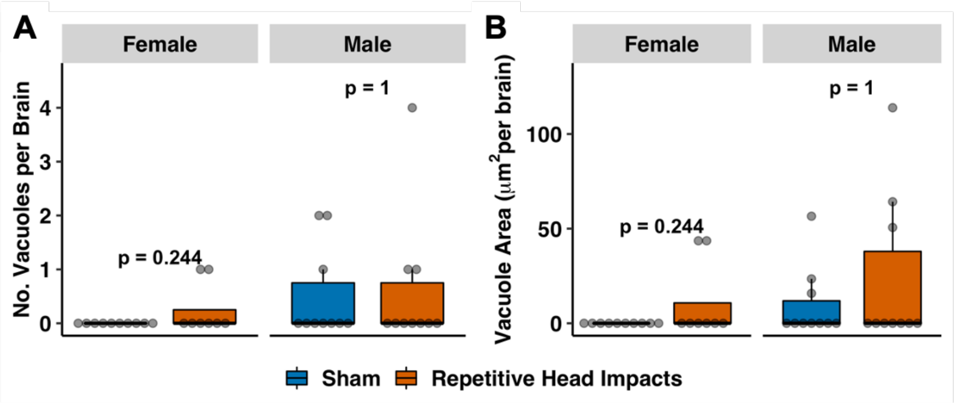


**Repetitive head impacts result in no acute neurodegeneration.** Repetitive head impacts elicit no acute (1.5h post-injury) neurodegeneration, neither seen as an **(a)** increased number of vacuoles and **(b)** vacuole area per brain. Boxplots contain individually plotted values with whiskers corresponding to the maximum 1.5 interquartile range. Within sex differences between sham and repetitive head impact conditions were analyzed with the Mann–Whitney *U* test with Bonferroni correction.

### Figure S6

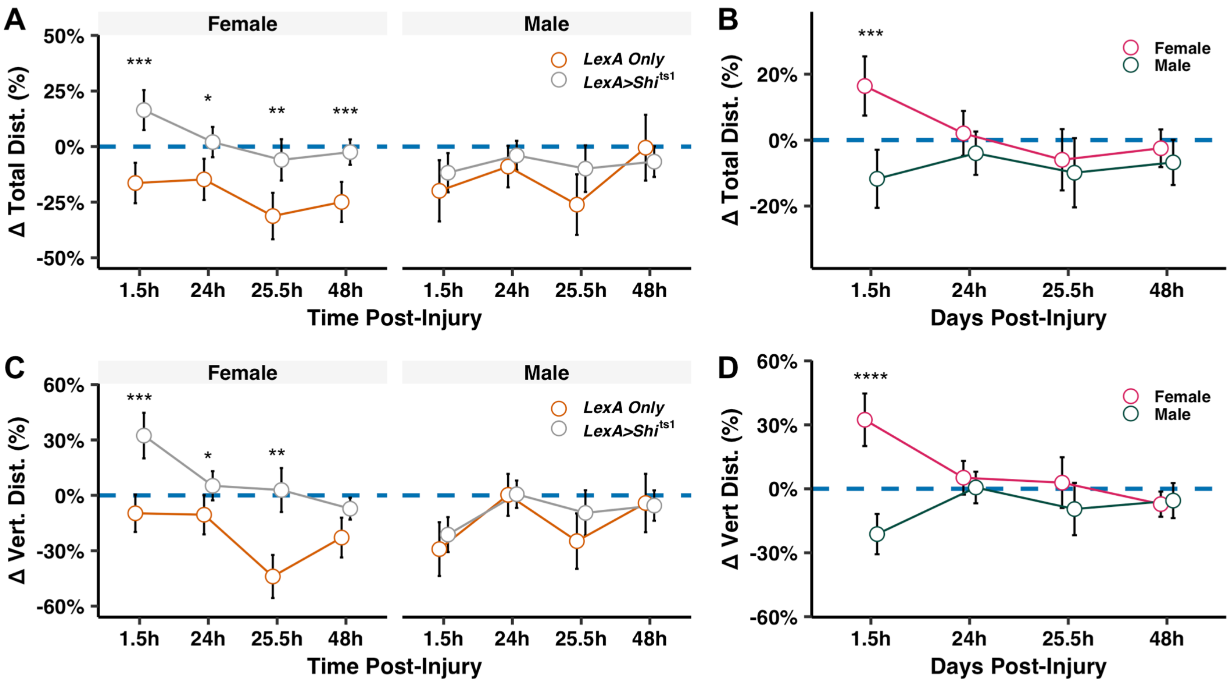


**Suppressing acute injury-induced neuronal activity following repetitive head impacts preferentially benefits females.** Blocking activity protects against acute climbing deficits in female flies, specifically **(a&b)** relative total distance and **(c&d)** relative vertical distance traversed. Plotted values are relative median values (compared to respective genotype sham) with 95% confidence interval error bars. Differences in relative climbing behavior were analyzed using the Mann–Whitney *U* test with Holm correction, between injured *LexA Only* and *Shi*$^{ts1}$-containing flies. *p<0.05, **p<0.01, ***p<0.001.

**Figure S7**

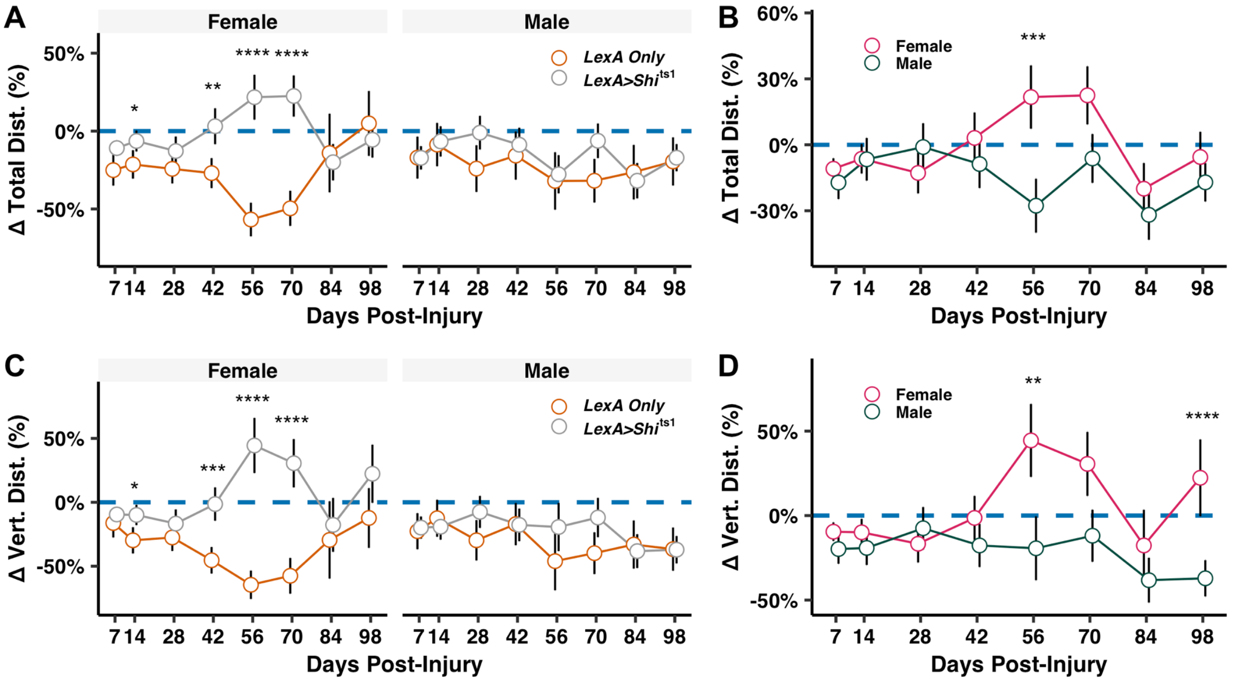


**Suppressing acute injury-induced neuronal activity following repetitive head impacts preferentially benefits females.** Blocking activity protects against chronic climbing deficits in female flies, specifically **(a&b)** relative total distance and **(c&d)** relative vertical distance traversed. Plotted values are relative median values (compared to respective genotype sham) with 95% confidence interval error bars. Differences in relative climbing behavior were analyzed using the Mann–Whitney *U* test with Holm correction, between injured *LexA Only* and *Shi*$^{ts1}$-containing flies. *p<0.05, **p<0.01, ***p<0.001.
